# Music Affects State Anxiety and Brain Connectivity

**DOI:** 10.1101/2023.05.18.541357

**Authors:** Mangor Pedersen, Daniel Shepherd, Geet Vashista, Amy Kercher, Michael. J. Hautus

## Abstract

Stress, anxiety, and depressive symptoms can be reduced by listening to music, but the underlying mechanisms remain unclear. To address this gap, we measured brain connectivity while participants listened to songs of different genres: ambient, pop, and metal. Additionally, affective ratings were obtained while participants (*n* = 30) listened to the six different songs, and subjective ratings of state anxiety were solicited at the terminus of each song. Electroencephalography (EEG) connectivity combining weighted Phase Lag Index and graph theory was utilised to document brain activity during listening. Repeated-measures ANOVA indicated that listening to more pleasant and less arousing songs was associated with lower self-reported state anxiety levels than songs rated unpleasant and highly arousing. Of interest, EEG alpha connectivity differed across two ambient songs, particularly in the frontal lobes, despite being from the same genre and rated as highly pleasant and low in arousal. We also observed a sex effect on EEG results, where female participants (*n* = 18) displayed stronger connectivity than male participants (*n* = 12). Combined, these results suggest that ambient songs reduce state anxiety but have divergent brain responses, possibly reflecting the complex nature of music listening, including sensory processing, emotion and cognition.

## Introduction

In many species, audition plays an important role as an alerting system, giving notice of threats that are either out of view or approaching during sleep. Higher evolved species have further used sound to express the opportunity for reproduction and to demarcate dominance hierarchies. These sonic functions are usually reliant upon sound produced by the organism itself, for example, birdsong or the rubbing of the hind legs by crickets. Human beings have acquired sophisticated skills to make sound, including music, which involves using musical instruments and/or the human voice to produce sounds possessing melody, rhythm, and harmony. Functionally, music can project aspects of an individual’s social identity (Shepherd & Sigg, 2015). Beyond social structures, music can provide enjoyment and pleasure to an individual, potentially enhancing their psychological well-being.

The use of music as a therapeutic agent has a short history relative to how long humans have produced music, and formally music therapy has taken the form of the creation of music, individually or in a group, or the consumption of pre-recorded music. It is this latter approach that, with the emergence over the last century of the phonograph, the vacuum tube, the transistor, and the integrated circuit, offers an individual the opportunity to manage emotion with sound. In contemporary times, the ubiquitous smartphone allows individuals to seek, acquire, and play music through loudspeaker units or earphones, typically at their convenience. The use of music can be used to boost self-esteem, reduce stress, anxiety, or depression, or regulate emotion (Harney et al., 2022; Saifman et al., 2023; Shepherd et al., 2023). Approaches to estimating the effectiveness of music to boost well-being can be divided into two gross categories: subjective (i.e., psychometric) responses, in which individuals self-refer, or objective measures (e.g., electrophysiological measurements). However, the scientific literature has been relatively subdued when presenting subjective or objective evidence relating to the self-administration of music therapy.

There are indications in the literature that music can be used as a self-administered sonic medicine to reduce anxiety (Harney et al., 2022). Anxiety is a significant and growing issue for people, with meta-analyses reporting that 31% of adolescents experienced elevated anxiety during the recent pandemic (Deng et al., 2023). Physiological symptoms, such as elevated heart and respiratory rates, increased sweating, shaking, agitation, and hypervigilance, are common (Lang et al., 2016). Psychological symptoms include excessive and difficult-to-control worry, rumination, catastrophic thinking, and negative biases, and while the focus of anxiety varies across individuals, symptoms can cause significant distress. Behaviourally, anxiety is associated with avoidance, with people commonly withdrawing from typically expected activities, causing marked interference in their lives. When feeling worried or distressed, individuals who use experiential avoidance and maladaptive coping show decreased emotional and psychological well-being (Fledderus et al., 2010). Interestingly, young people have been shown to use music maladaptively at times, with evidence that music choices can intensify pathological symptoms (Cheong & McFerran, 2016; Thomson et al., 2014).

Music has also been associated with decreased self-reported anxiety and sometimes with effects on blood pressure, cortisol and heart rate. However, psychophysiological effects are inconsistent and often not significantly associated with anxiety levels (Panteleeva et al., 2018). Without appropriate measures to regulate or address anxiety, it can cause marked distress and interference, contributing to comorbid psychological conditions such as depression, suicidal behaviour and substance misuse (Woodward & Fergusson, 2001). Current treatment guidelines recommend cognitive-behavioural therapy (CBT) and selective serotonin reuptake inhibitor (SSRI) medication and emphasise an urgent need for further interventions (Walter et al., 2020).

Finding innovative ways to support people with feelings of anxiety is a priority for clinicians and researchers. One avenue of investigation has been to look at brain communication during music listening. The necessity of multiple cortical area recruitment during music listening has led researchers to utilize connectivity and graph theory to study the brain (Bhattacharya & Petsche, 2005; Flores-Gutiérrez et al., 2007; Karmonik et al., 2016; Kay et al., 2012; Wu et al., 2012). These findings suggest that music-listening effects persist across multiple wave bands, notably alpha, and correspond with bilateral alterations in the frontotemporal EEG profile (Altenmüller et al., 2002). A suggestion has also been made that some level of lateralization across different cortical topographies within alpha band activity correlates with varying music patterns (Breitling et al., 1987; Schmidt & Trainor, 2001). Furthermore, unfamiliar music elicits a greater response than familiar music within cortical areas integral to emotional and music perception (Kumagai et al., 2017; Shahabi & Moghimi, 2016).

Building from this, only a few studies have explored the role of EEG alpha connectivity in anxiety specifically (e.g., Newson & Thiagarajan, 2019). Higher alpha connectivity correlates with anxiety and shyness in childhood and adolescence (Schmidt et al., 2022; Schmidt & Poole, 2021). In adults, alpha symmetry between the frontal lobes may be related to negative emotions (Demerdzieva & Pop-Jordanova, 2015). EEG alpha oscillatory activity is pertinent to audio processing (Lehtelä et al., 1997; Weisz et al., 2011). Wu et al. (2012) quantified alpha EEG connectivity while listening to music and found more efficient network characteristics during music listening than random noise. Others report stronger alpha wave connectivity and more efficient network topology occurs during music listening compared to eyes open and closed conditions (Zheng et al., 2018).

By better understanding the potential role of music in alleviating psychological distress and inducing parasympathetic nervous system dominance, the current study aims to offer a novel option to assist people to self-manage mild anxiety symptoms and reduce feelings of distress before they escalate. To this end, the present study will utilise six songs differing in affective character while obtaining a 64-channel EEG and conducting advanced functional connectivity analyses. The study hypothesizes that songs low in pleasantness and high in arousal will be associated with higher state anxiety. Given the limited EEG connectivity studies in the music literature, we treated the EEG analysis as exploratory.

## Materials and Methods

### Participants

Twelve males (*M_age_* = 23.3; *SD* = 2.14 years) and 18 females (*M_age_* = 23.1; *SD* = 1.60 years) served as volunteers for this study: Of these, 27 reported they were right-handed, while all participants reported normal hearing and no history of neurological disorders. Participants provided informed consent before data collection, and the study was performed with approval from the Auckland University of Technology Ethics Committee (AUTEC: 23/34).

### Psychometric measures

This study utilized two versions of the State-Trait Anxiety Inventory (STAI: (Spielberger et al., 1999). Firstly, the 40-item version divided equally into items measuring either state anxiety or trait anxiety was utilised as a general screen of participant anxiety. The 20-item State Anxiety Form requires participants to respond to several statements asking *how do you feel right now, at this moment,* using a four-point Likert Scale (1 = Not At All, 2 = Somewhat, 3 = Moderately So, 4 = Very Much So). The 20-item Trait Anxiety Form requires responses to statements asking participants how they *generally feel*, using a four-point Likert Scale (1 = Almost Never, 2 = Sometimes, 3 = Often, 4 = Almost Always). Secondly, the short-form version of the STAI, the STAI-6 (Marteau & Bekker, 1992), was utilised to measure State Anxiety following the presentation of each song. The STAI-6 used the same 4-point Likert scale as the 20-item State Anxiety Inventory and contained the six items that most correlate with the full 20-item scale.

The three dimensions of the self-assessment affective manikin were employed to gauge the participant’s emotional responses to the songs: pleasure, arousal and dominance (Betella & Verschure, 2016). As the music played, the participants were asked: *What do you think of this song? Please update the ratings if your opinion changes.* Here, a temporal dynamics approach was adopted, with participants rating the pleasantness (Extremely Unpleasant to Extremely Pleasant), excitability (Extremely Calming to Extremely Arousing), and focus (Extremely Distracting to Free to Think) of each song in real-time on a 1 to 10 scale. Additionally, a hedonic measure was presented. *I like this song*, rated on a scale from 0 (0% likability) to 4 (100% likability). All affective and hedonic measurements were obtained at 500-ms intervals while a song played.

### Songs

Six songs, each lasting 250 seconds, were selected based on their fit to the study’s objectives. These songs were specifically chosen based on their different affective characteristics (two ambient songs –Bagels and Weightless–, two heavy and fast songs –B.Y.O.B and 5 minutes Alone–, one complex mid-tempo song –Lateralus– and one pop song –Shape of You–), allowing for a comprehensive investigation into behavioural and neurophysiological changes during songs. Information about the six songs is provided in Supplementary Materials 1.

### Procedure

Participants were instructed not to consume caffeine before their session. Upon arrival, participants were provided information about the experiment protocol and equipment, and signed a consent form. The full STAI was completed before the EEG cap and insert headphones (E-A-RTONE with 3A inserts) were fitted.

The study used a single-subject design (Figure 1). All data were obtained in a commercially available sound-attenuating chamber equipped with a computer monitor and mouse, allowing participants to respond to the affective scales and the STAI-6. Following set-up, there was a two-minute preparation period during which the investigators ensured that all measurement equipment was functioning as desired. Then the first song was randomly played to the participant, who had been instructed to stay as still as possible while responding to the hedonic and affective scales via a computer mouse controlled by their dominant hand. At each song’s end was a four-minute quiet interval, with the first minute containing the STAI-6. For each participant, the six songs were randomised across the session, and once recording was initiated, a session took approximately 50 minutes.

**Figure 1:**
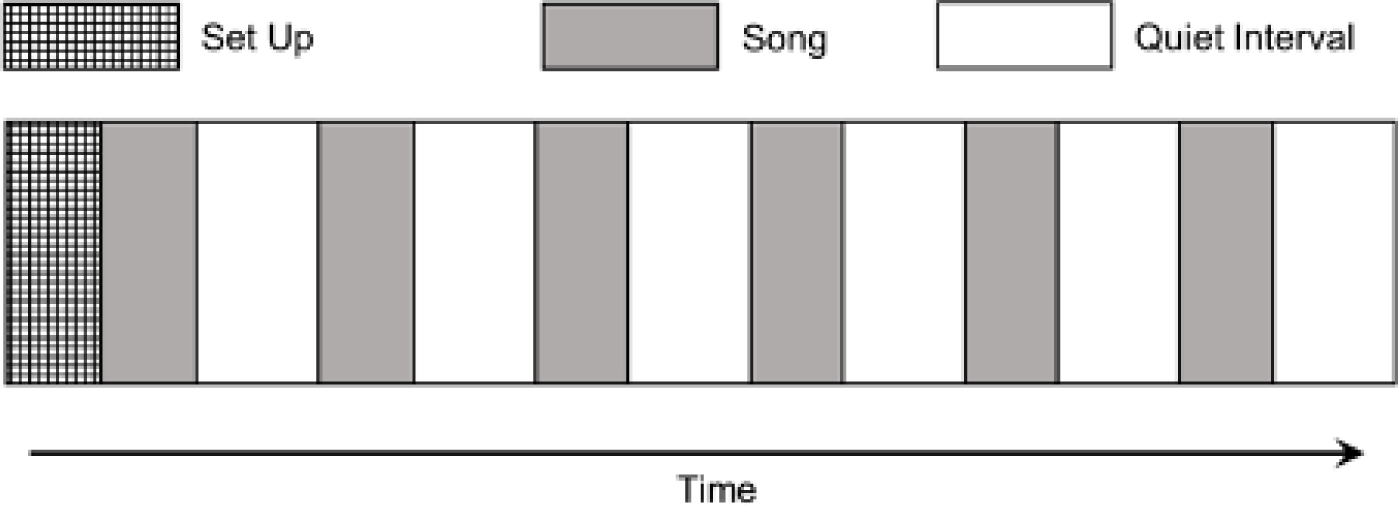
Temporal schematic of the procedure. The six songs were randomized across the session, while the first minute of each quiet interval required completion of the STAI-6.

### EEG data acquisition and pre-processing

EEG datasets were obtained with a 64-channel Quik-Cap Neo Net cap and SynAmps 2/RT amplifiers (Figure 2). The data was processed in a Brain Imaging Data Structure (BIDS) format (Gorgolewski et al., 2016) within the Python-based MNE software (version 1.3.1 – Gramfort et al., 2014). EEG was recorded from 64 electrodes, down-sampling to 250 Hz from an initial sampling rate of 1000 Hz. High-frequency signals were removed using a low-pass filter with a cut-off frequency of 100 Hz. 50 Hz line noise was eliminated using a notch filter. EEG data was virtually referenced using an average electrode referencing scheme. Noisy electrodes were interpolated with a spherical spline interpolation (Perrin et al., 1989).

**Figure 2:**
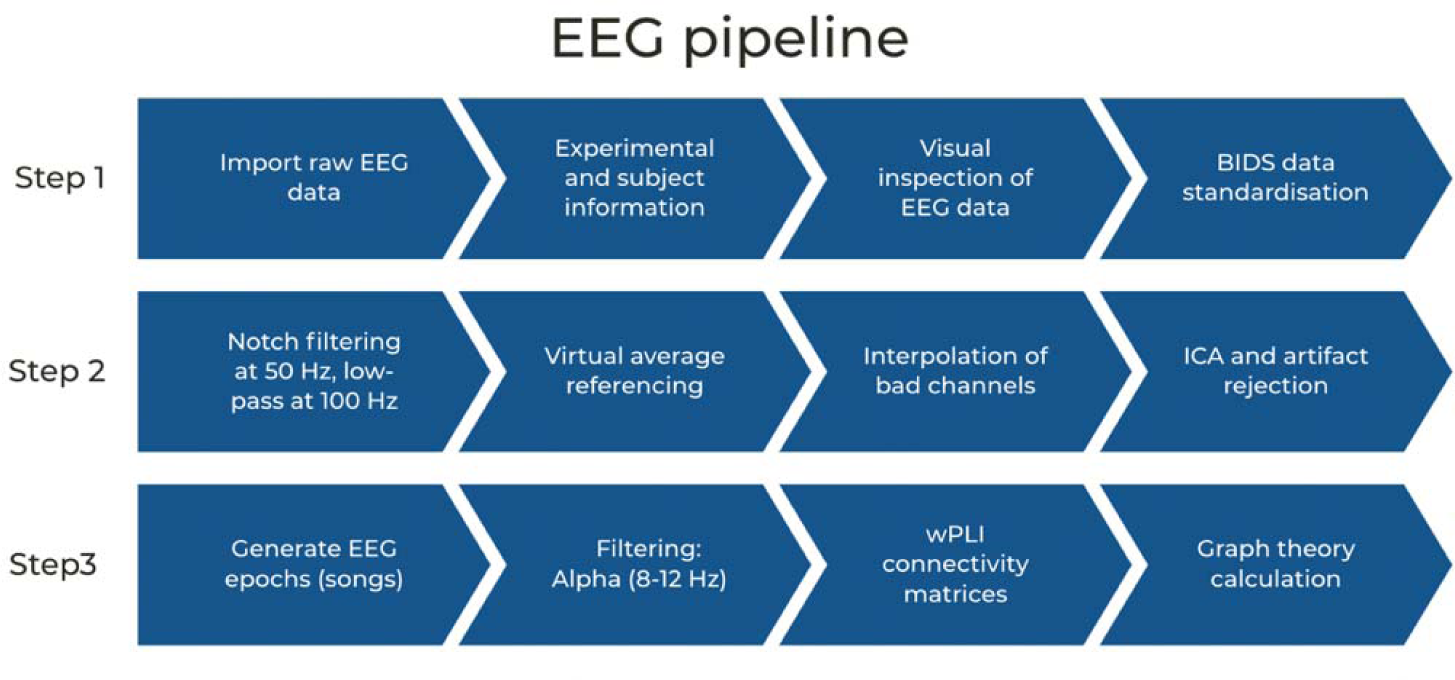
An outline of the EEG pre-processing pipeline used in the current study.

Independent Components Analysis (ICA: Urigüen & Garcia-Zapirain, 2015) was employed to remove artifacts from the EEG signals by decomposing the EEG signals into 20 independent components. Each component was examined visually, and those related to eye movements, muscle activity, or other sources of artifacts were eliminated. For each participant, the EEG data was converted into six epochs (one for each song) that were all 250 seconds long, resulting in 62,501 EEG time points per epoch.

56 electrodes were used for the connectivity: FP1, FPZ, FP2, AF3, AF4, F7, F5, F3, F1, Fz, F2, F4, F6, F8, FC5, FC3, FC1, FCz, FC2, FC4, FC6, T7, C5, C3, C1, CZ, C2, C4, C6, T8, TP7, CP5, CP3, CP1, CPZ, CP2, CP4, CP6, TP8, P7, P5, P3, P1, PZ, P2, P4, P6, P8, PO7, PO3, POZ, PO4, PO8, O1, Oz, O2. 8 electrodes were excluded for analysis as they were not overlying the brain (F11, F12, FT11, FT12, M1, M2, CB1, CB2). EEG data processing and analysis were conducted using an iMac 3.6 GHz 10-Core Intel Core i9 and 64GB RAM. A cumulative 8 hours of computation was done with MNE in Python (Gramfort et al., 2014). *EEG alpha connectivity*

Next, functional EEG connectivity was analysed using the weighted Phase Lag Index (wPLI) for each epoch/song (Vinck et al., 2011). wPLI is a measure of lagged synchronicity between pair-wise electrodes, denoting the phase differences of EEG signals between each electrode pair using the Hilbert transform (Bedrosian, 1963). The weights of the phase differences in each predefined bin are added to a histogram representing all phase differences, calculated by the imaginary magnitude of the cross-spectral density.

The *mne_connectivity.spectral_connectivity_time* function in MNE was used to calculate wPLI using a multitaper spectrum estimation of 7 Discrete Prolate Spheroidal Sequences (DPSS) taper windows (Slepian & Pollak, 1961). The cross-spectral density is calculated for each DPSS taper before summarising it to a weighted average. The normalized weighted phase synchronization index (wPLI) is calculated as the sum of the weighted phase differences divided by the sum of the weighted range of 0 to 1, with values approaching 1 indicating strong phase synchronization between signals (Stam et al., 2007; Vinck et al., 2011). wPLI was calculated within the alpha (8-12 Hz) frequency band. This study used four brain connectivity measures from fully connected, symmetric, and un-thresholded wPLI matrices. wPLI was calculated separately for each song, resulting in a matrix tensor of 64×64×6 (electrode *x* electrode *x* songs) for each participant. Three graph theory metrics provided EEG network properties:

i) *Node strength* is the total number of nodes (in this context, a *node* refers to EEG electrodes) and estimates a node’s connectedness to every other node in the network. Node strength is a metric for assessing a network’s general connectedness (Rubinov & Sporns, 2010).
ii) *Local efficiency* measures connectivity between neighbouring nodes in the network, calculated as the average of all pairs of a node’s neighbours’ inverse shortest path lengths. This metric measures communication effectiveness within a node’s local network (Latora & Marchiori, 2001).
iii) *Betweenness centrality* is defined as the number of shortest paths that run through a node in the network. A high betweenness centrality of a node suggests it is importance in a network with a capacity to relay information with other nodes. Such nodes are commonly referred to as hub nodes (Freeman, 1978)

### Statistical analysis

A repeated measures ANOVA was used to test for EEG and anxiety differences across the six songs. A 5 × 6 repeated measures ANOVA was used to test for differences in affective ratings, with the additional factor Time representing the average affective rating across five 50-second bins.

## Results

### Pleasantness ratings

Pleasantness ratings (Figure 3, top) indicate that the Bagels song was perceived as the most pleasant, followed by Shape of You and Weightless. A repeated-measures ANOVA containing two factors, song (six levels) and time (five levels), revealed a main effect of song (*F*(5,145) = 62.08, *p* < .001) and time (*F*(4,112) = 3.73, *p* = .496) on the pleasantness ratings. Additionally, a significant song-by-time interaction effect was also noted (*F*(20,560) = 72.08, *p* < .001). For the song variable, Bonferroni-adjusted testing revealed that the Bagels, Shape of You, and Weightless songs were rated significantly more pleasant than the 5 Minutes Alone, Lateralus, and B.Y.O.B songs (all *p* < .001). Additionally, the Lateralus song was rated significantly more pleasant than 5 Minutes Alone or B.Y.O.B (*p* < .001). Turning to the other main effect, time, *post hoc* testing revealed no significant differences once Bonferroni corrections were applied. Considering the significant interaction effect and the lack of cross-over in Figure 3 (top). It is unlikely that the main effect of the songs on pleasantness ratings is dependent upon time, as no significant main effect was obtained from the time factor.

**Figure 3:**
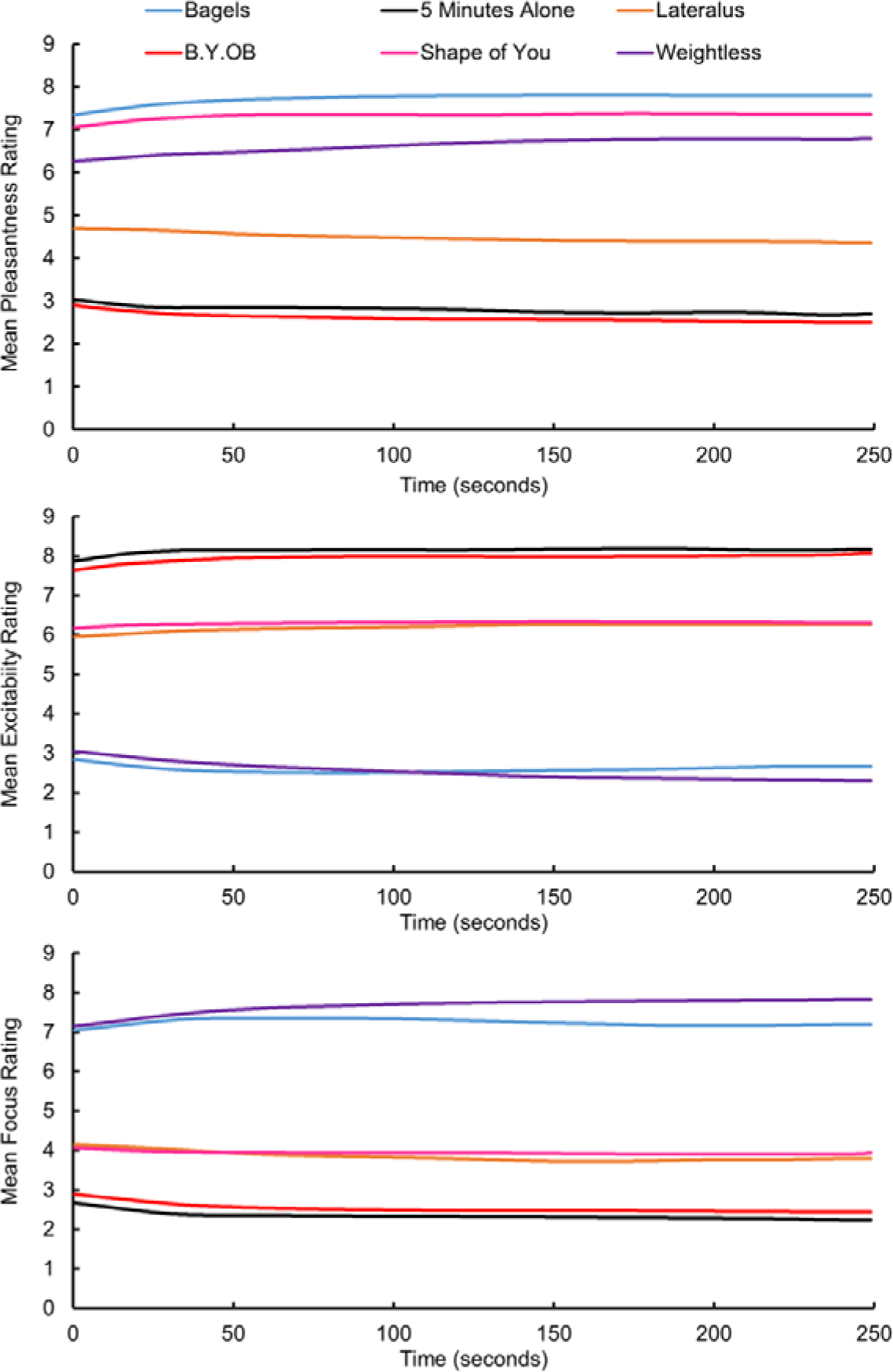
This figure displays mean affective scores pooled across the 30 participants as a function of time for the six songs. Affective ratings were measured at half-second intervals while the song played.

### Arousal ratings

When rating excitability (Figure 3, middle), participants rated 5 Minutes Alone and B.Y.O.B as the most arousing and the Bagels and Weightless track as least arousing. As for pleasantness, there was a main effect of songs (F(145) = 118.50, *p* < .001) and time (*F*(4,116) = 2.56, *p* = .042) on arousal, and again a song-by-time interaction (*F*(20,580) = 36.42, *p* = .496). For the six songs, all pair-wise comparisons were significant (*p* < .001), with the following exceptions being rated as having similar excitability: Bagels and Weightless, 5 Minutes alone and B.Y.O.B, and Lateralus and Shape of You. As for pleasantness, the marginal main effect noted for the time factor was not significant after the Bonferroni correction.

### Focus ratings

Focus ratings (Figure 3, bottom), indicating how dominant the song was in directing the participant’s attention, indicated that the two ambient songs (Bagels and Weightless) directed attention less and afforded greater focus than the other songs. There were main effects of song (*F*(5,145) = 71.48, *p* < .001) and time (*F*(4,116) = 3.19, *p* = .016) upon focus ratings. Significantly higher focus ratings, indicating a greater opportunity for free thought, were obtained between the Bagels and Weightless and all four songs (all *p* < .001). In contrast, the Lateralus and Shape of You songs were rated significantly higher than the 5 Minutes Alone and B.Y.O.B songs (all *p* < .05). For the time factor, the 3^rd^ bin (i.e., between 100 and 150 seconds) was associated with significantly higher focus ratings than the first (*p* = .020) and second epochs (*p* = .022).

### Hedonic ratings

Considering the hedonic ratings (Supplementary Materials 2), the Shape of You track was the most liked, followed by Bagels, with 5 Minutes Alone and B.Y.O.B being the least liked. There were main effects of song (*F*(5,145) = 57.62, *p* < .001) and time (*F*(4,116) = 3.89, *p* = .005) upon focus ratings, alongside a significant interaction effect (*F*(4,116) = 29.59, *p* <.001). Regarding the main effect of the songs, there were no statistical differences between any parings of the Bagels, Shape of You, and Weightless songs, nor between 5 Minutes Alone and B.Y.O.B. All other relationships were statistically significant (*p* < .005) across songs. Time, however, revealed no effect on hedonics following *post hoc* testing. To emphasize the individual differences in the hedonic data, a battery of Mann Whitney U tests reveals no sex differences for Bagels (*U* = 107.50, *p* = .983), 5 minutes alone (*U* = 80.50, *p* = .244), Shape of You (*U* = 105.00, *p* = .917), or Weightless (*U* = 80.50, *p* = .249) ratings. Hedonic ratings for Lateralus (*U* = 52.00, *p* = .017) and B.Y.O.B (*U* = 53.50, *p* = .019) were statistically higher for males than females (Supplementary Materials 3).

### Anxiety ratings

The total mean score for the full STAI was 38.9 ± 10.6 SD. The mean for the State anxiety subscale was 42.7 ± 9.9 SD (min = 26; max = 70) and the mean for the Trait anxiety subscale was 35.2 ± 10.2 SD (min = 21; max = 69). Figure 4 displays the STAI-6 scores obtained at the immediate terminus of each song. As higher scores indicate greater feelings of anxiety, it is evident that 5 Minutes Alone and B.Y.O.B were associated with greater feelings of anxiety, and Bagels and Weightless were associated with the least anxiety. A repeated-measures ANOVA revealed significant differences of anxiety ratings between the six songs (*F*(5,145) = 41.3, *p* < .001) between all pairings bar the following: Weightless and both Bagels (*p* = .927) and Shape of You (*p* = .311), and between 5 Minutes Alone and B.Y.O.B (*p* = .840).

**Figure 4:**
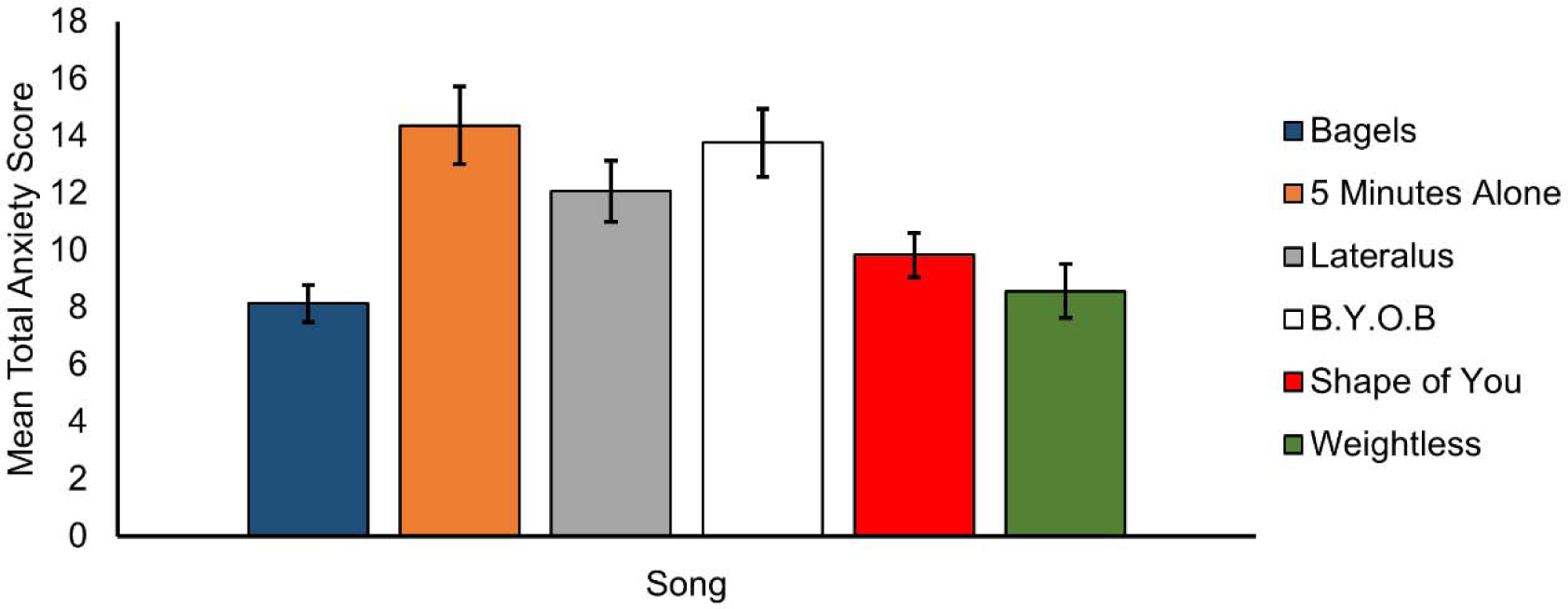
Mean STAI-6 scores for the six songs. Error bars = 95% confidence intervals.

### EEG alpha connectivity – descriptive results

EEG alpha connectivity analysis shows that Bagels had the lowest level of connectivity (average wPLI across all connections was 0.14 ± 0.12 SD), followed by B.Y.O.B (average wPLI across all connections was 0.16 ± 0.14 SD), Lateralus (average wPLI across all connections was 0.18 ± 0.16 SD), 5 Minutes Alone (average wPLI across all connections was 0.17 ± 0.14 SD), and Shape of You (average wPLI across all connections was 0.19 ± 0.16 SD) and Weightless: (average wPLI across all connections was 0.19 ± 0.16 SD). These values are derived from the average wPLI connectivity from the matrices in Figure 5A. Despite the difference in connectivity between songs seen in Figure 5A, we observed no difference in connectivity patterns between songs. All song pairs had Spearman’s *rho* correlation coefficients >0.99 (*p* < .001), suggesting that the topological patterns of the brain network do not change between songs (see Figure 5B). In other words, it is likely the strength of connectivity, not the organisation of connectivity, that changes during music listening. However, female participants (*n* = 18) displayed stronger connectivity than male participants (*n = 12*), particularly during Shape of You and Weightless. Male participants have the strongest connectivity during Lateralus. This observation suggests there may be a sex difference in connectivity during music listening (see Supplementary Materials 4).

**Figure 5:**
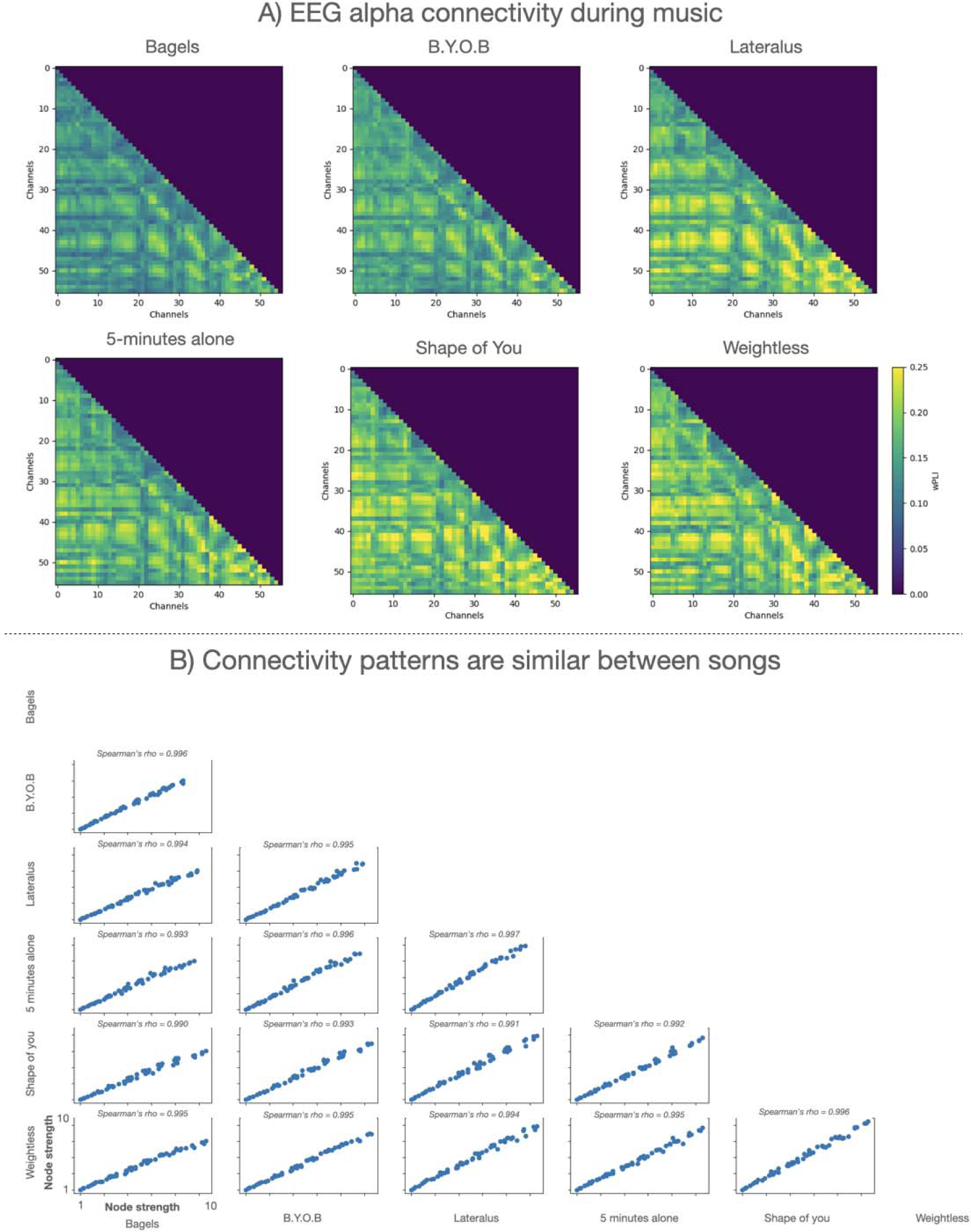
A) Connectivity matrices for all six songs. Colorbar is the same for all songs; connectivity weights are represented as wPLI. B) scatter-plots displaying the connectivity relationship between all songs – the blue dots represent the sum of connectivity (i.e., node strength) from the matrices in A, and each dot represents an EEG electrode averaged across subjects. Spearman’s correlation between variables is the number on top of each scatter plot.

### EEG alpha network topology as measured with graph theory

Next, a repeated measures ANOVA was performed to quantify the statistical difference in network topology between the six songs. Node strength (“overall connectivity of nodes”) and local efficiency (“local connectedness of nodes”), but not betweenness centrality (“nodal network hubs”), had several EEG electrodes with a significant ANOVA after correcting for multiple comparisons (False Discovery Rate at p <.05; Benjamini & Hochberg, 1995). The significant electrodes were predominantly located in the frontal lobe for node strength and fronto-temporo-parietal lobes for local efficiency (see a complete overview of significant EEG electrodes in Table 1). The largest effect size between songs was observed between the two ambient songs with reduced EEG alpha connectivity in the Bagels song compared to Weightless (see Figure 6).

**Figure 6:**
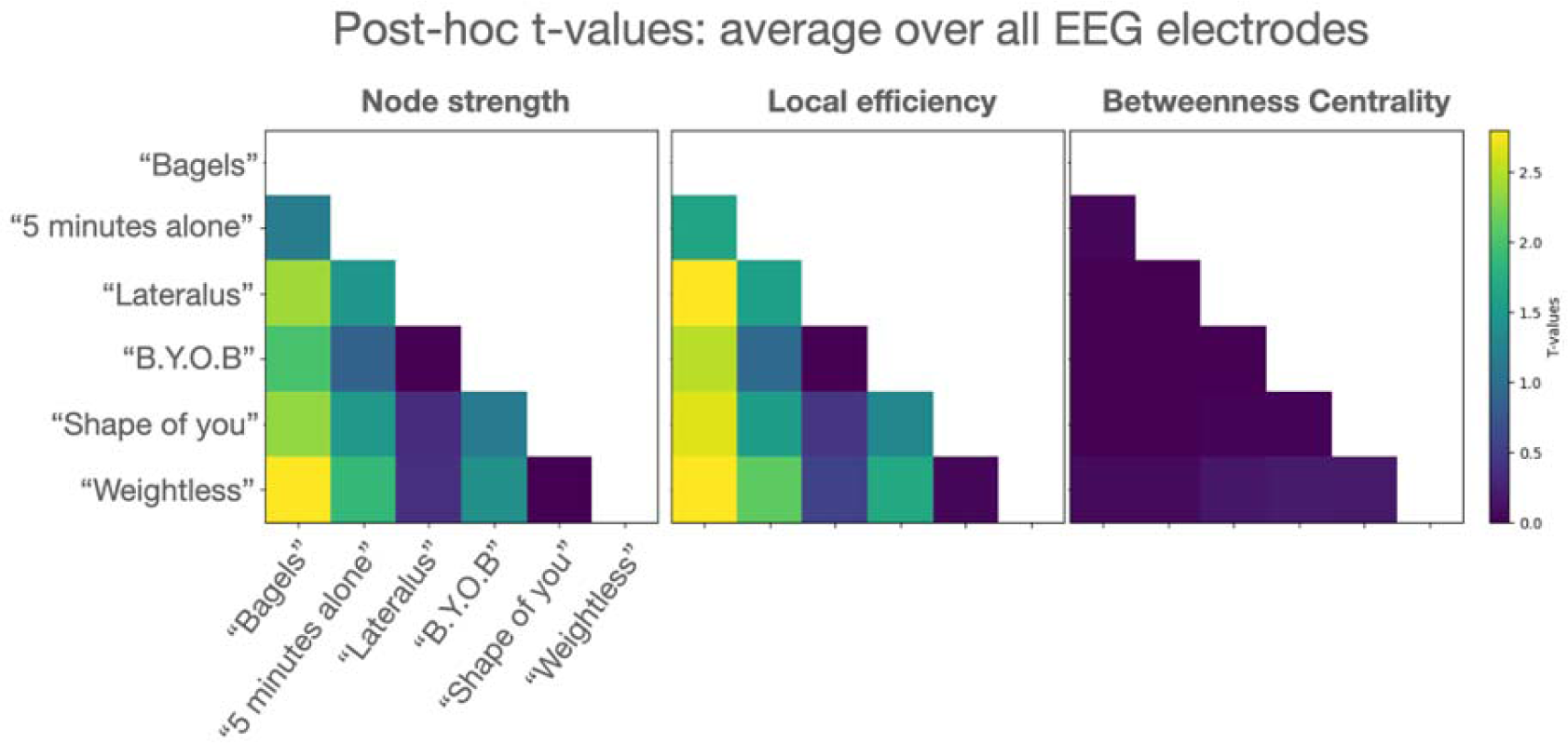
Example of pair-wise comparisons between songs – Left: the average t-values averaged across all electrodes for node strength. Middle: the average t-values averaged across all electrodes for Local efficiency. Right: the average t-values averaged across all electrodes for Betweenness centrality.

**Table 1.** Repeated measures ANOVA for EEG alpha connectivity metrics across 6 songs, corrected for multiple comparisons with False Discovery Rate (FDR). Note: No significant electrodes for Betweenness Centrality.

| Node strength |  | Local efficiency |  |
| --- | --- | --- | --- |
| FP1 | $F(5,145) = 5.45, p = .001$ | FP1 | $F(5,145) = 5.52, p = .002$ |
| FPZ | $F(5,145) = 5.56, p = .002$ | FPZ | $F(5,145) = 5.73, p = .001$ |
| FP2 | $F(5,145) = 5.14, p = .002$ | FP2 | $F(5,145) = 5.21, p = .005$ |
| AF3 | $F(5,145) = 4.48, p = .004$ | AF3 | $F(5,145) = 4.51, p = .005$ |
| AF4 | $F(5,145) = 4.78, p = .005$ | AF4 | $F(5,145) = 4.84, p = .005$ |
| F7 | $F(5,145) = 5.57, p < 0.001$ | F7 | $F(5,145) = 5.12, p = .001$ |
| F4 | $F(5,145) = 4.52, p = .007$ | F4 | $F(5,145) = 4.44, p = .007$ |
| F6 | $F(5,145) = 4.31, p = .009$ | F9 | $F(5,145) = 4.23, p = .009$ |
| F8 | $F(5,145) = 4.06, p = .009$ | FC6 | $F(5,145) = 4.05, p = .007$ |
| FC6 | $F(5,145) = 4.46, p = .005$ | TP7 | $F(5,145) = 4.88, p = .003$ |
| T7 | $F(5,145) = 3.81, p = .009$ | CP5 | $F(5,145) = 4.64, p = .002$ |
| TP7 | $F(5,145) = 4.21, p = .005$ | CP3 | $F(5,145) = 4.30, p = .006$ |
| TP8 | $F(5,145) = 3.83, p = .008$ | TP8 | $F(5,145) = 4.12, p = .009$ |
| P5 | $F(5,145) = 3.84, p = .008$ | P5 | $F(5, 145) = 5.73, p < 0.001$ |
| | | P3 | $F(5,145) = 4.96, p = .002$ |
| | | P6 | $F(5,145) = 4.40, p = .007$ |
| | | P8 | $F(5,145) = 4.06, p = .007$ |
| | | PO3 | $F(5,145) = 4.11, p = .006$ |
| | | O1 | $F(5,145) = 4.53, p = .003$ |

The ANOVA analyses were performed a second time with sex as a covariate, given the descriptive sex differences in connectivity discussed in the previous paragraph and shown in Supplementary Materials 4. No EEG electrodes were statistically significant after False Discovery Rate Correction, suggesting that sex differences are important in our results.

### EEG post-hoc investigation

Conducting *post-hoc* comparisons was not statistically appropriate for the two significant ANOVAs (node strength and local efficiency), given that 6 songs and 64 electrodes would result in 1920 tests. However, the whole-brain averaged *t*-test matrices in Figure 5 suggest that the largest pair-wise effect size between songs was between Bagels and the other songs. In light of the ANOVA model described in the previous paragraph, this result is likely to be influenced by the brain connectivity patterns in the female participants.

## Discussion

The study’s findings suggest that listening to more pleasant and less arousing music was associated with lower self-reported state anxiety levels than listening to songs rated as unpleasant and arousing. Moreover, the ambient song, Bagels, had reduced EEG alpha connectivity compared to several other songs, particularly in female participants. These findings show that pleasant and less arousing music (e.g., ambient music) can be linked to lower state anxiety and is associated with complex neurobiological responses across sexes, likely reflecting the multimodal characteristics of music listening.

The adaptive use of music in stress management shows promise, and music provides a potentially easy, appealing and cost-efficient anxiolytic that people can administer wherever they are (McFerran et al., 2018). Our study suggests that music can assist people in reducing subjective feelings of anxiety, and associated changes in brain connectivity. Using music may assist people to self-manage mild-moderate distress. The current findings reinforce suggestions from Thomson et al. (2014), arguing that careful music creation and selection are key. Importantly, selecting music that has a calming effect and is appealing to listeners appears essential. The current research also supports previous evidence suggesting that some styles of music increase arousal and feelings of anxiety (Cheong & McFerran, 2016; Thomson et al., 2014). This finding is consistent with previous literature, for example, when waiting for a medical appointment (Yeoh & Spence, 2023).

In the current study, the participants reported that ambient music was the most pleasant (Bagels) and was associated with the lowest state anxiety scores (Bagels and Weightless). In contrast, heavy and fast songs (5 minutes alone and B.Y.O.B) were associated with higher anxiety. This finding extends previous work examining the differential use of musical genres by young people, in which venting negative emotion through music was associated with higher levels of depression, using music for distraction (avoidance) predicted anxiety and stress, and choosing music for entertainment and positive affect was associated with low depression (Thomson et al., 2014). Many playlists, artists and apps (e.g., https://www.calm.com/ and https://www.ipnos.com/) aim to reduce anxiety and induce relaxation through meditation, music and ambient noise; however, few have been empirically tested, and fewer have undergone neuroscientific testing. This novel approach in the current study extends previous research on musical styles and demonstrates shifts in anxious arousal associated with ambient music.

Although the EEG alpha activity literature is extensive and often with conflicting results (Miljevic et al., 2023), EEG alpha connectivity findings are still nascent. Despite this, previous research has shown that EEG alpha activity and connectivity may increase during music listening (Flores-Gutiérrez et al., 2009; Petsche et al., 1997; Wu et al., 2012). Of note, the two songs with the lowest state anxiety ratings – the ambient songs, Bagels and Weightless – also had the most divergent EEG alpha connectivity patterns. Music listening affects multiple complex processes in the brain, including sensory perception/integration, cognition and emotion (Krumhansl, 2002). In this context, it is unlikely that the EEG findings solely relate to a reduction in anxiety. These findings are similar to previous fMRI work, which found bilateral and decreased activity with people listening to pleasant music (Koelsch et al., 2006). This activity, predominantly appearing in the limbic system, Heschl’s gyrus (auditory cortex), and inferior frontal structures, may be a pattern consistent with the present study’s finding in the ambient song Bagels.

The disparity in connectivity strength between male and female participants was a principal finding. Sex differences between brain connectivity profiles have been documented (Gong et al., 2011). Ingalhalikar et al. (2014) found that female brains have greater interhemispheric connectivity and global network integration, whereas male participants have greater intrahemispheric connectivity and modularity. These brain-specific differences may partly explain the current findings showing sex-specific connectivity profiles to music. Surprisingly, the ambient song (Weightless) had divergent connectivity strength between the sexes, while the other ambient song (Bagels) had similar connectivity profiles between the sexes. These connectivity patterns that differed between sexes were overlying frontal and parietal lobes (Table 1). These are brain areas typically associated with higher-order cognitive processing rather than the introspective default mode network in the brain’s (posterior) midline (Raichle, 2015). This, in turn, suggests that our observations reflect brain changes related to cognition. This ties in with theories of alpha activity that suggest that the neurobiological mechanism of alpha activity is related to cortical inhibition, an essential mechanism for cognitive processes (Klimesch, 2012). As EEG alpha connectivity is related to cognitive processing, the ambient song Weightless may appear more complex, with unconventional musical patterns, resulting in increased EEG connectivity.

When interpreting the study’s findings, several caveats should be considered. A limitation of this study is the sample size. However, regarding the central limit theorem and the within-groups design used in the current study, a sample of 30 participants is typically deemed sufficient in EEG research. Secondly, the demographics of the participants. All were between 18 and 25 and in good health, so the current findings cannot readily be generalised to a clinical or the general population with any degree of confidence. Future research would benefit from replicating the current results in source space, targeting the specific brain regions that change connectivity to different music styles.

## Conclusion

Our findings show that songs deemed unpleasant and arousing were associated with higher levels of self-reported state anxiety than songs rated as pleasant and less arousing. We also show diverging EEG alpha connectivity between songs of the same genre and being regarded as highly pleasant and low in arousal, potentially reflecting sex differences in the music listening experience, which involves complex sensory, emotional, and cognitive processes.

