## Supplementary Materials for "Music Affects State Anxiety and Brain Connectivity"

Supplementary materials #1:

Information of the 6 songs used in the current study.

1. Bagels, by New Zealand artist Benee (Genre: Ambient). A slow instrumental punctuated by a short narrative of the singer having a challenging day and searching foretheir happy place to revitalise themselves.
2. 5 minutes alone, by Pantera (Genre: Groove Metal). A heavy, swinging, hard metal song characterised by an aggressive narrative in which the singer fantasies about attacking one of his concert attendees.
3. Lateralus, by Tool (Genre: Progressive Metal). It is a heavy but slow tempo song with its time signatures manifesting the Fibonacci sequence. The lyrics are cryptic, but can be loosely interpreted as the singer taking a (lateral) journey to facilitate self-understanding.
4. B.Y.O.B, by System of a Down (Genre: Alternative Metal). An aggressive and fast anti-war song in which the singer beseechs, “why do we always send the poor?”.
5. Shape of You, by Ed Sheeran (Genre: Pop). A light dance number describes the development of an interpersonal relationship between two individuals.
6. Weightless, by Marconi Union (Genre: Ambient). Described as the “worlds most relaxing song” by Time Magazine (Grossman, 2011), Weightless was developed in collaboration with the British Academy of Sound Therapy to induce relaxation.

*Supplementary Materials #2:*

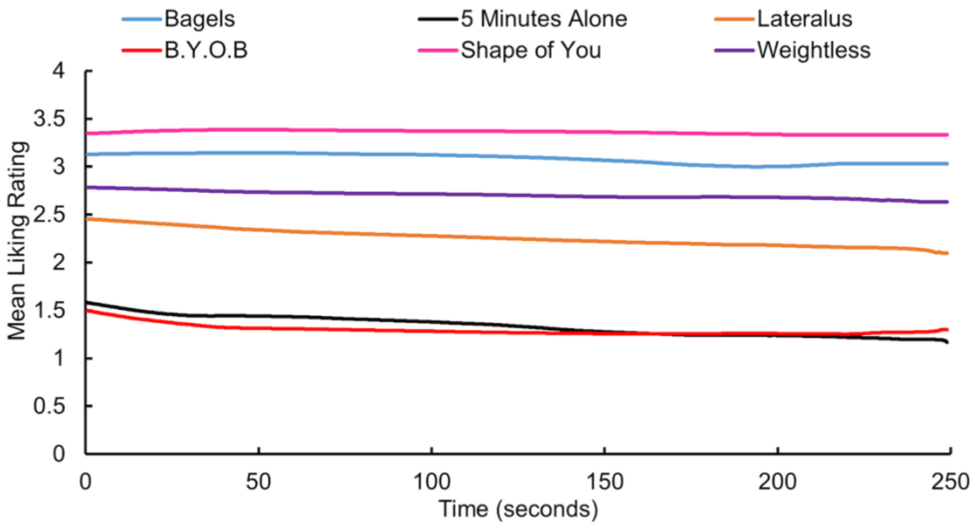


*This figure displays mean hedonic scores, over time.*

*Supplementary materials #3: Mean hedonic ratings for each participants as a function of song for females (top-half of table) and males (bottom half). Parentheses contain standard deviations.*

| | | Song Artist | | | | | |
|---|---|---|---|---|---|---|---|
| Females | Age | Bagels | 5 minutes alone | Lateralus | B.Y.O.B | Sheeran | Weightless |
| 1 | 24 | 3.24 (0.66) | 3.08 (0.58) | 3.00 (0.06) | 0.23 (0.53) | 1.98 (0.75) | 3 (0.00) |
| 2 | 24 | 3.00 (0.00) | 0.23 (0.91) | 1.18 (1.07) | 0.12 (0.60) | 2.05 (0.21) | 2.14 (0.35) |
| 3 | 24 | 2.70 (0.46) | 3.72 (0.59) | 3.00 (0.00) | 1.93 (0.84) | 2.31 (0.46) | 2.49 (0.5) |
| 4 | 24 | 3.89 (0.24) | 0.69 (0.60) | 1.48 (0.87) | 0.33 (0.85) | 4.76 (0.56) | 4.17 (0.66) |
| 5 | 23 | 2.31 (0.47) | 1.91 (0.71) | 2.26 (0.44) | 1.34 (0.75) | 3.02 (0.14) | 2.19 (0.39) |
| 6 | 24 | 3.84 (0.38) | 0.35 (0.96) | 2.17 (0.38) | 0.82 (0.81) | 3.90 (0.34) | 3.70 (0.46) |
| 7 | 19 | 4.28 (0.96) | 2.54 (0.50) | 3.00 (0.00) | 2.15 (0.36) | 3.00 (0.06) | 3.75 (0.43) |
| 8 | 23 | 3.93 (0.25) | 0.26 (0.84) | 1.18 (0.55) | 0.16 (0.67) | 3.87 (0.33) | 4.83 (0.56) |
| 9 | 24 | 3.58 (0.49) | 0.58 (0.85) | 1.34 (0.76) | 0.45 (1.07) | 3.91 (0.29) | 2.16 (0.47) |
| 10 | 23 | 4.75 (0.58) | 2.11 (0.31) | 2.98 (0.13) | 0.29 (0.76) | 3.00 (0.06) | 3.56 (0.50) |
| 11 | 24 | 2.12 (0.33) | 1.21 (0.60) | 1.41 (0.66) | 0.08 (0.48) | 3.00 (0.00) | 2.09 (0.28) |
| 12 | 24 | 1.22 (0.63) | 0.16 (0.67) | 0.90 (0.98) | 3.76 (0.43) | 3.92 (0.28) | 2.12 (0.32) |
| 13 | 24 | 2.99 (0.20) | 1.18 (0.61) | 1.54 (0.78) | 0.16 (0.68) | 3.84 (0.37) | 3.00 (0.00) |
| 14 | 24 | 3.76 (.043) | 0.81 (0.93) | 1.88 (0.78) | 1.58 (0.76) | 2.13 (0.34) | 3.88 (0.34) |
| 15 | 19 | 3.01 (0.12) | 3.00 (0.00) | 3.00 (0.00) | 2.24 (0.43) | 3.95 (0.22) | 3.00 (0.00) |
| 16 | 24 | 3.01 (0.10) | 0.75 (0.81) | 3.00 (0.00) | 1.40 (0.74) | 3.92 (0.27) | 2.19 (0.40) |
| 17 | 23 | 2.62 (0.64) | 1.07 (0.73) | 3.00 (0.00) | 1.16 (0.48) | 3.85 (0.36) | 3.00 (0.04) |
| 18 | 22 | 1.86 (0.62) | 1.71 (0.70) | 0.58 (1.18) | 0.63 (1.03) | 2.82 (0.39) | 2.18 (0.40) |
| Mean | 23.1 (1.61) | 3.13 (0.90) | 1.48 (1.17) | 2.05 (0.87) | 1.09 (0.97) | 3.33 (0.87) | 3.01 (0.82) |
| Males | Age | Bagels | 5 minutes alone | Lateralus | B.Y.O.B | Sheeran | Weightless |
| 1 | 25 | 2.24 (0.43) | 1.08 (0.42) | 2.99 (0.11) | 1.09 (0.42) | 3.94 (0.23) | 3.02 (0.13) |
| 2 | 25 | 3.87 (0.34) | 2.43 (0.71) | 4.15 (0.97) | 2.48 (0.50) | 3.94 (0.24) | 2.09 (0.33) |
| 3 | 24 | 2.35 (0.5) | 3.60 (0.49) | 2.29 (0.57) | 1.44 (0.82) | 3.01 (0.17) | 2.23 (0.63) |
| 4 | 18 | 3.89 (0.31) | 1.26 (1.18) | 2.47 (0.52) | 0.28 (0.85) | 3.00 (0.00) | 3.00 (0.06) |
| 5 | 23 | 2.98 (0.15) | 2.11 (0.33) | 2.98 (0.13) | 3.00 (0.00) | 3.02 (0.13) | 1.58 (0.88) |
| 6 | 24 | 2.81 (0.39) | 1.09 (0.42) | 3.00 (0.00) | 1.30 (0.72) | 3.80 (0.41) | 2.28 (0.96) |
| 7 | 25 | 2.33 (0.42) | 2.70 (0.84) | 3.05 (0.22) | 2.10 (0.44) | 3.86 (0.35) | 2.20 (0.45) |
| 8 | 24 | 3.96 (0.22) | 2.05 (0.21) | 2.98 (0.13) | 2.96 (0.22) | 3.00 (0.00) | 3.56 (0.50) |
| 9 | 24 | 4.21 (0.97) | 0.39 (1.01) | 2.05 (0.56) | 0.57 (0.93) | 3.56 (0.50) | 3.00 (0.00) |
| 10 | 23 | 2.16 (0.91) | 1.35 (0.69) | 2.77 (0.58) | 2.18 (0.39) | 2.43 (0.50) | 0.47 (1.07) |
| 11 | 20 | 3.54 (0.50) | 1.66 (0.81) | 4.53 (0.82) | 4.90 (0.42) | 3.74 (0.44) | 2.19 (0.39) |
| 12 | 24 | 3.44 (0.56) | 2.31 (0.52) | 3.47 (0.51) | 3.76 (0.43) | 3.83 (0.40) | 3.61 (0.84) |
| Mean | 23.25 (2.14) | 3.15 (0.76) | 1.84 (0.87) | 3.06 (0.71) | 2.17 (1.35) | 3.43 (0.51) | 2.44 (0.88) |

*Supplementary materials #4:*

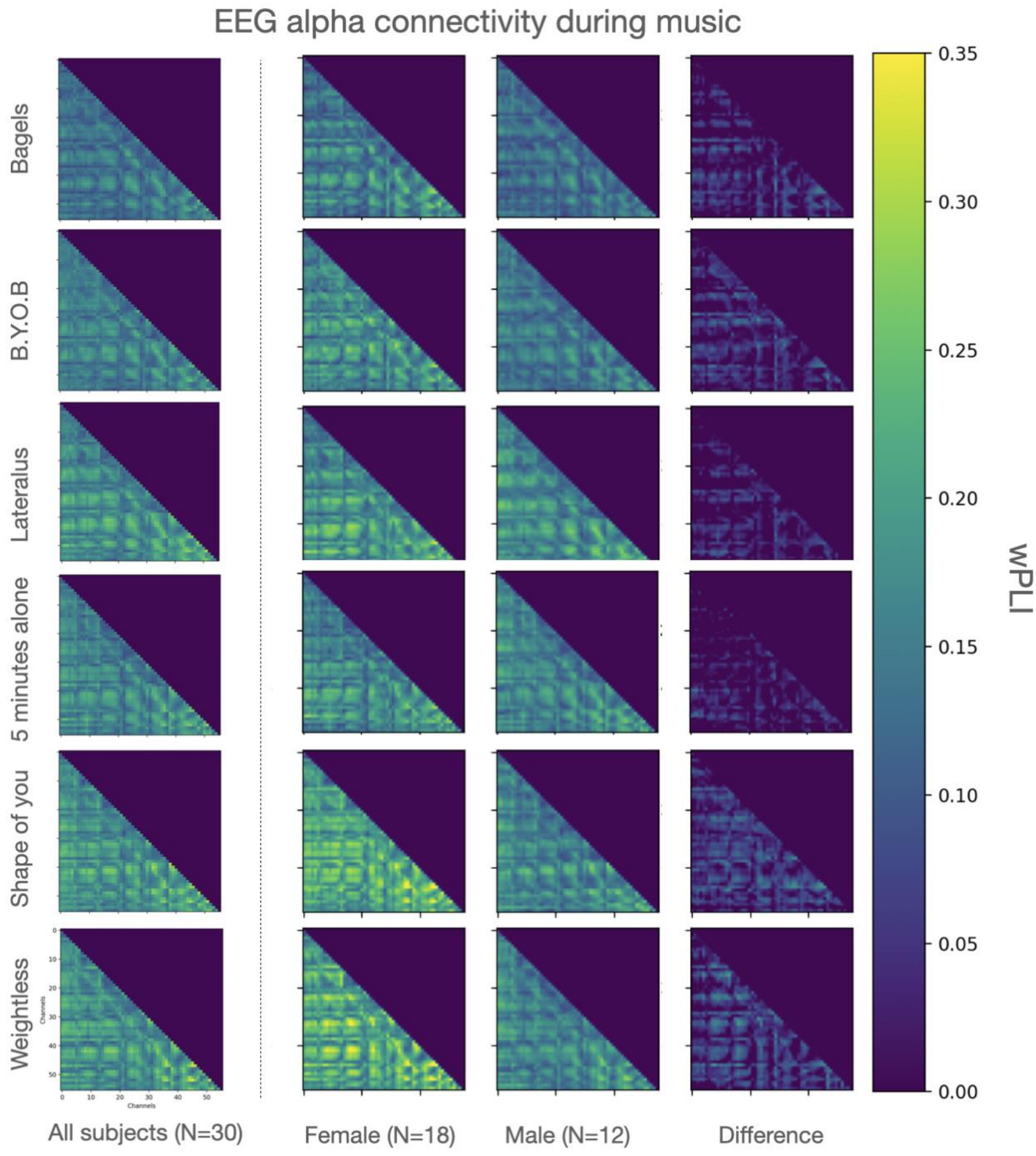
